# DEG/ENaC/ASIC channels vary in their sensitivity to anti-hypertensive and non-steroidal anti-inflammatory drugs

**DOI:** 10.1101/2020.02.12.946681

**Authors:** S. Fechner, I. D’Alessandro, L. Wang, C. Tower, L. Tao, M.B. Goodman

## Abstract

The degenerin channels, epithelial sodium channels, and acid-sensing ion channels (DEG/ENaC/ASICs) play important roles in sensing mechanical stimuli, regulating salt homeostasis, and responding to acidification in the nervous system. They have two transmembrane domains separated by a large extracellular domain and are believed to assemble as homomeric or heteromeric trimers. Based on studies of selected family members, these channels are assumed to form non-voltage gated and sodium-selective channels sensitive to the anti-hypertensive drug, amiloride. They are also emerging as a target of nonsteroidal anti-inflammatory drugs (NSAIDs). *C. elegans* has more than two dozen genes encoding DEG/ENaC/ASIC subunits, providing an excellent opportunity to examine variations in drug sensitivity. Here, we analyze a subset of the *C. elegans* DEG/ENaC/ASIC proteins to test the hypothesis that individual family members vary not only in their ability to form homomeric channels, but also in their drug sensitivity. We selected five *C. elegans* DEG/ENaC/ASICs (DEGT-1, DEL-1, UNC-8, MEC-10 and MEC-4) that are co-expressed in mechanosensory neurons and expressed gain-of-function *‘d’* mutant isoforms in *Xenopus laevis* oocytes. We found that only DEGT-1d, UNC-8d, and MEC-4d formed homomeric channels and that, unlike MEC-4d and UNC-8d, DEGT-1d channels were insensitive to amiloride and its analogs. As reported for rat ASIC1a, NSAIDs inhibit DEGT-1d and UNC-8d channels. Unexpectedly, MEC-4d was strongly potentiated by NSAIDs, an effect that was decreased by mutations in the putative NSAID binding site in the extracellular domain. Collectively, these findings reveal that not all DEG/ENaC/ASIC channels are amiloride-sensitive and that NSAIDs can both inhibit and potentiate these channels.

**Summary:** Animal physiology depends on degenerin, epithelial sodium, and acid-sensing ion channels (DEG/ENaC/ASICs). By measuring the sensitivity of three *C. elegans* DEG/ENaC/ASICs to five amiloride analogs and five NSAIDs, we show that individual channels have distinct pharmacological footprints.

## Introduction

The degenerin, epithelial sodium, and acid-sensing ion channels (DEG/ENaC/ASICs) are present in most, if not all metazoan genomes and expressed in diverse tissues, including the epithelia of several organs and in the central and peripheral nervous systems (Eastwood and Goodman, 2012; Kellenberger and Schild, 2002). At least two DEG proteins are known to be mechanosensitive, ENaCs are constitutively active and can be regulated by shear stress, and ASICs are activated by proton binding (Eastwood and Goodman, 2012). The DEGs were identified in *C. elegans* by virtue of their role in mechanosensation and by gain-of-function mutations that cause neuronal degeneration (Chalfie and Wolinsky, 1990; Driscoll and Chalfie, 1991; Huang and Chalfie, 1994). The ENaCs were identified *via* expression of rodent cRNAs in Xenopus oocytes followed by functional screening (Canessa et al., 1995). The proteins that form acid-sensing ion channels (ASICs) were based on their homology to DEGs and ENaCs (Waldmann et al., 1997)(García-Añoveros et al., 1997; Kellenberger and Schild, 2002).

All of these proteins have short intracellular amino and carboxy terminals and two transmembrane domains linked by a large extracellular domain which is divided into structures described by a hand holding a ball: wrist, finger, ball, and knuckle. This view has emerged from high-resolution structures derived from x-ray diffraction of protein crystals (Baconguis et al., 2014; Dawson et al., 2012; Gonzales et al., 2009; Jasti et al., 2007; Noreng et al., 2018) and cryo-electron microscopy of chicken ASIC1a (Sun et al., 2018; Yoder et al., 2018) and human ENaC (Noreng et al., 2018). Individual proteins assemble into trimers and form a pore along a common three-fold access at the center of the complex. The extracellular finger domain exhibits more sequence variation than other domains and the general topology of all family members are assumed to share the same fold as ASIC1a. Although ENaCs are formed from three distinct proteins, many DEGs and ASICs can form both homomeric and heteromeric channels. Thus, the ensemble of functional channels is expanded not only by genetic variation of individual channel subunits, but also by the formation of heteromeric channels.

The ENaC channels are crucial for salt homeostasis and are blocked by amiloride (Garty and Benos, 1988; Palmer, 1992), a classic anti-hypertension drug that functions as an open-channel blocker (Schild et al., 1997). Sensitivity to amiloride and its analogs is not limited to mammalian family members or to ENaCs, however, but is also seen in DEG/ENaC/ASIC channels expressed in invertebrates. Amiloride is a potent (sub-micomolar *IC*_50_) blocker of DEG and ENaC channels, but is at least hundred-fold less potent as a blocker of ASIC channels (Canessa et al., 1994; Goodman et al., 2002; Vullo and Kellenberger, 2019). Despite widespread findings of amiloride as an inhibitor, amiloride is also reported to potentiate the activity of two DEG/ENaC/ASIC channels (Adams et al., 1999; Elkhatib et al., 2019). It is not known whether amiloride inhibition and potentiation arise from binding to similar or distinct sites. Among channels inhibited by amiloride, variations in sensitivity could reflect differences in binding affinity or in the efficacy of inhibition. It remains unknown whether or not sensitivity to amiloride or its analogs is a universal feature of DEG/ENaC/ASIC channels.

The ASIC channels are implicated in neurological disease and in pain sensation, but there are no potent and selective small molecule inhibitors of ASICs available (Boscardin et al., 2016; Hanukoglu and Hanukoglu, 2016; Kellenberger and Schild, 2015; Vullo and Kellenberger, 2019). Evidence is emerging that ASICs are targets of non-steroidal anti-inflammatory drugs (NSAIDs). In particular, ibuprofen is an effective (micromolar *IC*_50_) allosteric inhibitor of H^+^-evoked ASIC1a currents and mutations in the wrist and the first transmembrane domain (TM1) reduce the apparent affinity for ibuprofen (Lynagh et al., 2017). This finding implicates NSAIDs as an additional class of small molecules affecting the function of DEG/ENaC/ASIC channels.

Whereas mammalian genomes have nine genes encoding ENaC and ASIC proteins (Hanukoglu and Hanukoglu, 2016; Kellenberger and Schild, 2002), the *C. elegans* genome harbors more than two dozen genes encoding DEG/ENaC/ASIC proteins (Goodman and Schwarz, 2003; Hobert, 2013). Thus, the *C. elegans* set of DEG/ENaC/ASIC proteins offers an excellent opportunity to examine variations in the biophysical properties within this superfamily. As an entry point for exploration, we expressed five DEG/ENaC/ASIC proteins in *Xenopus* oocytes individually and tested their ability to form functional channels, measured their permeability to monovalent cations, and examined their response to amiloride and its analogs as well as a set of NSAIDs. The five DEG/ENaC/ASIC proteins we studied, DEGT-1d, DEL-1d, UNC-8d, MEC-10d and MEC-4d, are expressed in touch receptor neurons [MEC-4, MEC-10, and DEGT-1 (M Chatzigeorgiou et al., 2010; Driscoll and Chalfie, 1991; Huang and Chalfie, 1994)], mechanical nociceptors [UNC-8, MEC-10, DEL-1 (Chatzigeorgiou et al., 2010; Tavernarakis et al., 1997)], and motor neurons [UNC-8, DEL-1 (Tavernarakis et al., 1997)]. All five proteins are either known or proposed to contribute to the formation of mechanosensitive ion channels (Chatzigeorgiou et al., 2010; Liu et al., 2020; O’Hagan et al., 2005; Tao et al., 2019).

We engineered these channels to be constitutively active based upon reported gain-of-function mutations that cause necrotic cell death – the “d” isoform. Here, we found that DEL-1d fails to generate any detectable current on its own and confirmed prior work showing that MEC-10d (Goodman et al., 2002) is also not sufficient to generate current. By contrast, DEGT-1d forms a homomeric channel that is insensitive to amiloride and its analogs, has a more negative reversal potential than other channels, and is blocked by NSAIDs. Both MEC-4d and UNC-8d formed channels blocked by amiloride and carried currents, consistent with prior work (Goodman et al., 2002; Wang et al., 2013). Unexpectedly, MEC-4d current was strongly potentiated by NSAIDs and sensitivity to these drugs was decreased by mutations in the extracellular domain that affect inhibition of ASIC1a by ibuprofen, a frontline NSAID drug. Collectively, these findings reveal that not all DEG/ENaC/ASIC channels are amiloride-sensitive and that NSAIDs can both potentiate and inhibit these channels.

## Materials and Methods

### Expression constructs and molecular biology

Plasmids carrying native cDNAs encoding MEC-4, MEC-10 and other *C. elegans* DEG/ENaC/ASIC proteins derived from the *C. elegans* genome are subject to deletions and recombination when propagated in standard bacterial strains (Chalfie et al., 2003; Goodman et al., 2002). Previously, we circumvented this outcome using SMC4, a bacterial strain specifically derived for this purpose (Chalfie et al., 2003; Goodman et al., 2002). Here, we used an alternative strategy that enables us to propagate expression plasmids in standard bacterial strains (NEB 5-alpha Competent *E. coli*, High Efficiency): synthetic cDNAs codon-optimized for expression in insect cells. Accordingly, we obtained plasmids containing synthesized codon-optimized cDNAs encoding full-length MEC-4, MEC-10 and DEGT-1 (GenScript) in the pGEM-HE oocyte expression vector (Liman et al., 1992). DEL-1 was codon-optimized for expression in *C. elegans* (IDT) based on the predicted sequence reported in wormbase release WS250. The predicted isoforms encoded by the *del-1* locus have been modified in a more recent database releases (WS274) and these changes are evident only in the amino terminal domains. WS274 predicts three isoforms and the isoform we used from WS250 encodes 18 amino acids that are not represented in the updated predictions. Unlike MEC-4, MEC-10, DEL-1 and DEGT-1, the UNC-8 isoform was not codon-optimized. This plasmid was obtained from L. Bianchi, used in prior studies (Matthewman et al., 2018, 2016; Miller-Fleming et al., 2016; Wang et al., 2013), and encodes the shortest of four predicted splice variants (R13A1.4d).

We studied the expressed channels as constitutively-active or degeneration (‘d’) isoforms based on gain-of-function mutations identified in forward genetic screens or engineered into homologous residues. Because co-expressing MEC-2 yields larger currents than expressing MEC-4d alone and co-expressing MEC-2 with UNC-8d is either indifferent or may yield to larger currents than expressing UNC-8d alone (Brown et al., 2007; Goodman et al., 2002; Matthewman et al., 2016), we co-expressed MEC-2 with all DEG/ENaC/ASIC channels studied here.

We used *in vitro* mutagenesis (Q-5 Site-directed Mutagenesis Kit, NEB) to introduce the mutations creating *d* isoforms using the following primers. All mutations were introduced into plasmids encoding wild-type protein, except the last two mutant isoforms that introduced second mutations into plasmids encoding MEC-4[A713T].

MEC-4[A713T]: CTCTTGACTGACTTCGGTGG, GTTCACGAAACCGTAGGCTT;
MEC-10[A676V]: AAGATGATGGTTGACTTCGGC, CACGATACCGTAGGCCTC
DEGT-1[A813T]: CTCTTGACTGAGATCGGAGG, CAGGAACAAGTTGTAGGAGC.
UNC-8[G347E]: AAAGATGCGGAAGCCATCACA, GAGGTCGCTCAATCCAAAAG
DEL-1[A603V]: AATTTGATGGTCGATATGGGAG, GAACCATGAATATGACTCG
MEC-4[A713T, E704K]: CTCTTGACTGACTTCGGTGG, GTTCACGAAACCGTAGGCTT (A713T); GGCCTACGGTTTCGTGAACCTCT, TTGGACTCGGTGAGCATCTCGAA (E704K)
MEC-4[A713T,E704A]: CTCTTGACTGACTTCGGTGG, GTTCACGAAACCGTAGGCTT (A713T); GCGGACTCGGTGAGCATCTC, AGCCTACGGTTTCGTGAACCTC (E704A)

### RNA preparation, validation and oocyte injection

For each channel isoform, we generated capped RNAs (cRNA) using *in vitro* transcription (mMESSAGE mMACHINE T7 kit, Ambion) and quantified cRNA concentration spectroscopically (NanoDrop2000, Thermo Fisher Scientific). We validated the size and integrity of cRNAs using gel electrophoresis (Reliant®RNA Gels, 1.25% SKG, Lonza). To each cRNA sample, we added buffer (2 μL 10x MOPS buffer, Lonza) and loading dye (8 μL, B0363S, NEB) and loaded denatured (70 °C, 10 minutes) samples alongside an RNA ladder (2-4 μL, ssRNA ladder, N0362S, NEB). The resulting gels were stained with ethidium bromide for 30 minutes (0.5 μg/mL ddH_2_0, Thermo Fisher Scientific), washed in ddH_2_0 (30 minutes), and visualized with UV light.

*Xenopus laevis* oocytes were isolated from gravid females (NASCO) modified from standard procedures (Liu and Liu, 2006). Briefly, frogs were anesthetized with MS-222 (0.5%, 1 hour), follicles were removed, opened with forceps and transferred to OR-2 solution. For defolliculation, oocytes were incubated twice (1st 45 min, 2nd variable) in OR-2 containing 3 mg/ml collagenase type IV (Sigma, C-9891). Defolliculated oocytes were stored in ND96 solution containing (in mM): NaCl 96, KCl 2.5, MgCl_2_ 1, CaCl_2_ 1.8, HEPES 5 at pH 7.6, adjusted with NaOH, containing 10 mg/ml Penicillin-streptomycin solution (Sigma, P0781). The OR-2 solution contained (in mM): NaCl 82.5, KCl 2.5, MgCl_2_ 1, HEPES 5 at pH 7.6. We injected cRNA encoding a single DEG/ENaC/ASIC isoform (5 ng) and MEC-2 cRNA (15 ng) in each oocyte. We reduced cRNA amounts to 3 ng (MEC-4d isoforms) and 9 ng (MEC-2) for cells used to collect ibuprofen dose-response curves. We maintained oocytes at 18°C in modified Leibovitz medium (L-15) (Sigma Aldrich) supplemented with gentamicin (144 μM) (Gibco) and amiloride (300 μM) for 2–9 days, as described (Brown et al., 2007). Oocytes expressing UNC-8d were maintained in the same solution with additional 100 μM benzamil. To minimize the impact of variation in expression efficiency and endogenous ion channels, we report data from oocytes derived from at least three donor frogs for each channel isoform.

### Whole-cell recordings and external solutions

Membrane current was measured by two-electrode voltage clamp (TEVC; OC-725C, Warner Instruments, LLC) at room temperature (21–24°C). Electrodes (~0.3 MΩ) were fabricated from borosilicate glass (G100TF-4, Warner Instruments, LLC) on a horizontal puller (P-97; Sutter Instruments) and filled with 3 M KCl. Analog signals (current, voltage) were digitized (Instrutech, ITC-16), filtered at 200 Hz (8-pole Bessel filter), and sampled at 1 kHz. A 60-Hz notch filter (FLA-01, Cygnus Technology, Inc.) was used to reduce line noise. This equipment was controlled by Patchmaster software (HEKA) on a windows PC.

Unless otherwise indicated, oocytes were superfused with control saline containing (in mM): Na-gluconate (100), KCl (2), MgCl_2_ (2), CaCl_2_ (1) and HEPES (10), adjusted to pH 7.4 with NaOH. Drugs were diluted from stock solutions and added to control saline. For pH 6.4 and 8.4 solution, we replaced HEPES in the control saline with 10 mM PIPES and 10 mM TAPS, respectively.

We purchased drugs from the indicated suppliers and established stock solutions in DMSO. We used stock solutions of 0.1 M, except for phenamil (0.01M) and for ibuprofen and aspirin dose-response experiments (0.24 M). Drugs were obtained from these suppliers: amiloride (amil, Sigma-Aldrich, A7410), benzamil (benz, Sigma-Aldrich, B2417), 5-(N-Ethyl-N-isopropyl)amiloride (EIPA, Sigma-Aldrich, A3085), phenamil (phen, Cayman Chemical, 14308), benzamidine (Fluka, 12072), ibuprofen (Ibu, Sigma-Aldrich, SLBR3566V), R-ibuprofen (R-ibu, Cayman Chemical, 16794), S-ibuprofen (S-ibu, Cayman Chemical, 375160), aspirin (Asp, Sigma-Aldrich, A2093), salicylic acid (SA, Sigma-Aldrich, S5922), diclofenac (Diclo, Sigma-Aldrich, D6899), flurbiprofen (Fibu, Sigma-Aldrich, F8514).

### Measuring current, drug sensitivity, and reversal potential

We used two voltage protocols to measure membrane current and its response to amiloride analogs and NSAIDs: a voltage ramp (from −100 to +100 mV in 1 s) and voltage steps (from −100 to +40 mV or +60 mV, in 20 mV increments). Both protocols included a conditioning step to −85 mV, which we used to measure current amplitude. In all cases, the holding potential was −60 mV. We applied these protocols repetitively during the application of drugs and solutions with modified ion composition; the ramp protocol to measure reversal potentials.

To determine which channels were indifferent, inhibited, or activated by each of the ten drugs we tested, we applied each drug a final concentration of 30 μM and measured the difference or drug-sensitive current, *I*_drug_, at −85mV. Average values were taken from epochs during step and ramp protocols when cells were held at −85 mV (Fig. 1 A, top) and averaged over three voltage presentations in the absence and presence of each drug. Next, we compared *I*_drug_ in experimental and control (uninjected) oocytes, computing the distribution of Δ*I*_drug_ using estimation statistics (Ho et al., 2019). We used an analogous strategy to assess which channels were indifferent, inhibited or activated by protons. Specifically, we measured *I*_ΔpH_ by subtracting the current measured at −85 mV in the presence of pH 6.4 saline from the current measured at pH 8.4, a 100-fold increase in H^+^ concentration. We compared *I*_ΔpH_ in experimental and control oocytes, computing Δ*I*_ΔpH_, We measured reversal potentials from voltage ramps (Fig. 1 A, top, −100 to +100 mV in 1 s). Drug dose-response curves were measured at voltage steps between −100 and either +40 or +60 mV in 20 mV increment steps. Three replicates of the voltage step protocol were averaged to derive current amplitude in the presence of each drug concentration.

**Figure 1:**
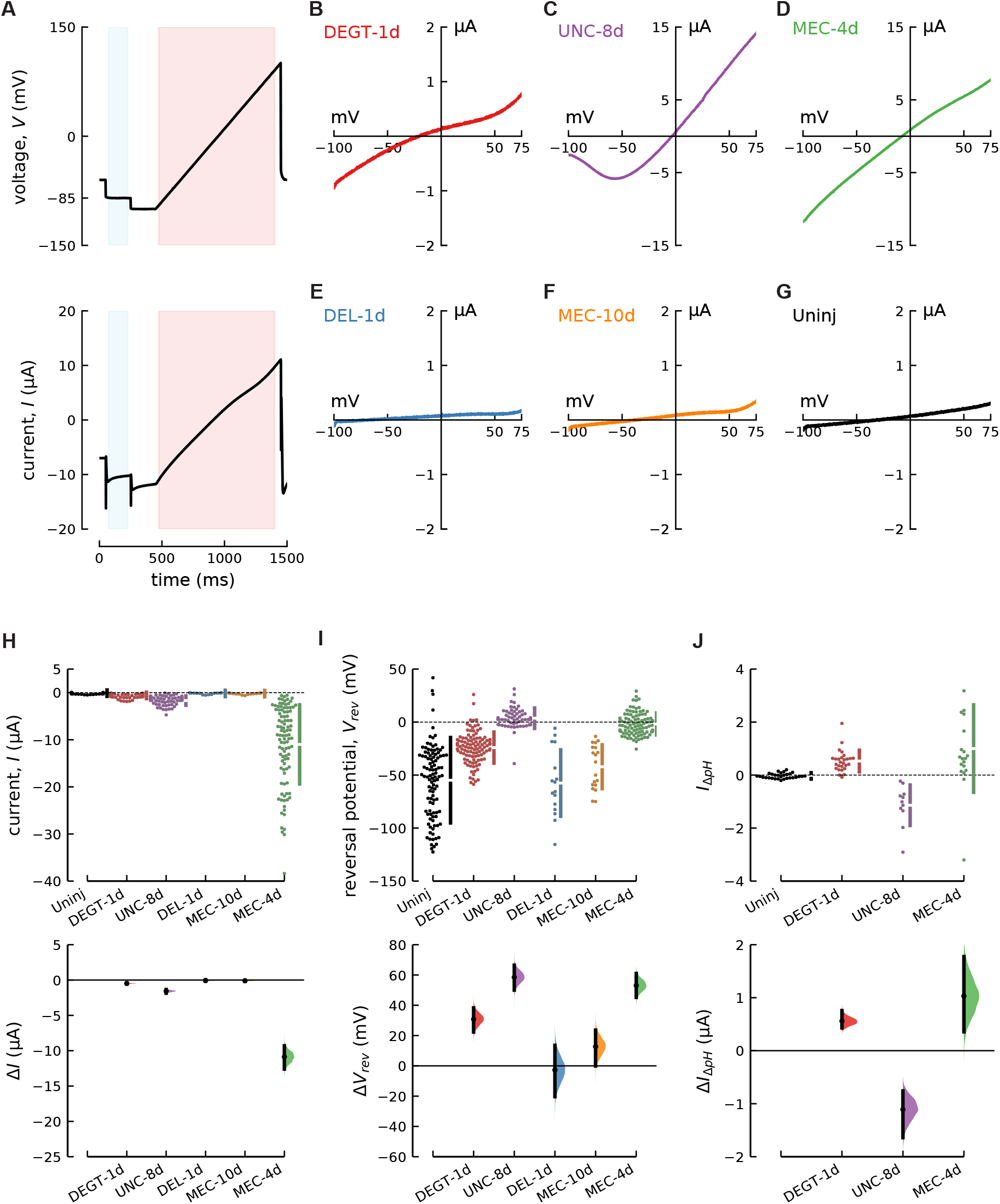
The DEG/ENaC/ASIC subunits DEGT-1d, UNC-8d and MEC-4d form homomeric channels in Xenopus oocytes, while DEL-1d and MEC-10d likely do not. **(A)** Graphical representation of the voltage-clamp protocol (top) and total current measured in an oocyte expressing MEC-4d (bottom). Shaded areas are epochs at a holding potential of −85mV (gray) and a voltage ramp from −100 to 100 mV (pink) used to measure current amplitude and reversal potential, respectively. **(B-G)** Current-voltage curves measured in oocytes co-expressing MEC-2 and DEGT-1d **(B)**, UNC-8d **(C)**, MEC-4d **(D)**, DEL-1d **(E)**, MEC-10d **(F)** and uninjected oocytes (**G**). (**H)** Total current, *I* (at −85 mV) as a function of uninjected and expressed proteins (top) and estimation plot (bottom) showing the effect of channel protein expression relative to uninjected oocytes (*ΔI*). **(I)** Reversal potential (*V*_rev_) as a function of uninjected and expressed proteins (top) and estimation plot (bottom) showing the effect of channel protein expression relative to uninjected oocytes (*ΔV*). Mean values, 95% confidence intervals, and statistical analyses are in Table 1. (**J)** Change in current (at −85 mV) induced by a pH shift from 8.4 to 6.4 (*IΔpH*) in uninjected oocytes and oocytes expressing DEGT-1d, UNC-8d, MEC-4d (top) and estimation plot (bottom) showing the effect size relative to uninjected oocytes (*ΔI*). The mean values, 95% confidence intervals, and statistical tests for the data in panels B-G are in Table 1.

### Data Analysis, curve fitting, and figure generation

Mean values and reversal potentials were calculated using MATLAB (R2014b) (https://github.com/wormsenseLab/AnalysisFunction.git). ANOVA (two-way) followed by multiple comparison with the Holm-Sidak method (P< 0.05) was performed in Sigma Plot 12.5. Estimation statistics were performed in Python using Data Analysis using Bootstrap-Coupled ESTimation or DABEST (Ho et al., 2019). Individual dose response curves were fitted with the Hill equation, with a Hill coefficient set to one, to calculate the *EC*_50_ = *I*_*max*_ * [*x*^*n*^/(*EC*_50_^*n*^ + *x*^*n*^)] and *IC*_50_ = *I*_*max*_ * *IC*_50_^*n*^/(*IC*_50_^*n*^ + *x*^*n*^) using IgorPro 6.37. We used the mean values for *EC*_50_ and *IC*_50_ to compute an average fit to the pooled and averaged data. Confidence intervals for mean relative permeability ratios were calculated in Python with DescrStatsW. Figures were prepared in python jupyter notebooks (https://github.com/wormsenseLab/JupyterNotebooksDEGENaCPharm.git).

### Sequence analysis

The following ion channel sequences were used to generate a phylogenetic tree (Fig. 7 A) and alignments (Fig. 7 C): ACD-1 (C24G7.2), ACD-2 (C24G7.4), ACD-3b (C27C12.5b), ACD-4 (F28A12.1), ACD-5 (T28F2.7), ASIC-1 (ZK770.1), ASIC-2 (T28F4.2), DEG-1 (C47C12.6.1), DEGT-1 (F25D1.4), DEL-1 (E02H4.1), DEL-2a (F58G6.6a), DEL-3 (F26A3.6), DEL-4 (T28B8.5), DEL-5 (F59F3.4), DEL-6 (T21C9.3), DEL-7 (C46A5.2), DEL-8 (C11E4.3), DEL-9 (C18B2.6), DEL-10 (T28D9.7), DELM-1 (F23B2.3), DELM-2 (C24G7.1), EGAS-1 (Y69H2.11), EGAS-2 (Y69H2.12), EGAS-3 (Y69H2.2), EGAS-4 (F55G1.13), FLR-1 (F02D10.5), MEC-4 (T01C8.7), MEC-10 (F16F9.5), UNC-8d (R13A1.4d), UNC-105e (C41C4.5e), rASIC-1 (NP_077068), rASIC-2 (Q62962.1), rASIC-3 (NP_775158.1), rASIC-4 (Q9JHS6.1), rαENaC (NP_113736.1), rβENaC (NP_036780.1), rγENaC (NP_058742.2), hδENaC (001123885.2). The alignment for calculating the phylogentic tree was generated with MUSCL and http://www.phylogeny.fr/ (Dereeper et al., 2010, 2008) and visualized with figtree v1.4.4 (Rambaud, 2018). The alignment was generated with Clustal Omega (Fig. 7 C).

### Online supplemental material

Fig. S1 shows how DEGT-1 is affected by amiloride ibuprofen concentrations higher than 30μM as well as how amiloride sensitivity is independent of the presence or absence of MEC-2. Fig. S2 shows the dose-dependence and voltage sensitivity of UNC-8d inhibition by benzamil and MEC-4d inhibition by EIPA.

## Results

We expressed five *C. elegans* DEG/ENaC/ASIC proteins in Xenopus oocytes to determine which could form homomeric ion channels. The proteins we studied, DEGT-1d, DEL-1d, UNC-8d, MEC-10d and MEC-4d, are expressed in three classes of mechanoreceptor neurons: touch receptor neurons or TRNs (ALM, PLM, AVM, PVM), and polymodal nociceptors (ASH, PVD) and stretch-sensitive motor neurons (D-MNs). To enhance constitutive currents and increase the likelihood of detecting DEG/ENaC/ASIC-dependent currents, we used mutant or *d* isoforms of each of these proteins linked to neuronal degeneration. We also co-expressed all channel subunits with MEC-2, based on prior studies showing that this yields larger currents than expressing MEC-4d alone (Brown et al., 2007; Goodman et al., 2002; Matthewman et al., 2016; Zhang et al., 2004). Individual proteins were deemed capable of forming homomeric ion channels if total membrane current (measured at −85 mV) and its reversal potential differed from those measured in uninjected, control oocytes. We sought additional evidence that each protein formed active channels by measuring sensitivity to amiloride and its analogs and to a panel of NSAIDs.

### DEGT-1d, but not DEL-1d forms homomeric channels

Previous research showed that the DEG/ENaC/ASIC subunits MEC-4d and UNC-8d, but not MEC-10d can form homomeric channels when exogenously expressed in *Xenopus laevis* oocytes (Goodman et al., 2002; Wang et al., 2013). Co-expression with MEC-2 increases MEC-4d (Brown et al., 2008; Goodman et al., 2002; Matthewman et al., 2016), but not UNC-8d currents (Matthewman et al., 2016). On average, oocytes expressing MEC-4d generated inward currents at −85 mV that were approximately 7-fold larger than those expressing UNC-8d and 17-fold larger than those expressing DEGT-1d. Oocytes expressing MEC-10d and DEL-1d, by contrast, generated currents that were indistinguishable from those recorded in uninjected oocytes (Fig. 1, A-I, Table 1). Consistent with prior reports that oocytes expressing DEG/ENaC/ASIC channels become sodium-loaded during the incubation period (Brown et al., 2007; Canessa et al., 1994; Goodman et al., 2002), membrane current reversed polarity near 0 mV in oocytes expressing MEC-4d and UNC-8d (Fig. 1, Panel C, D, and I, Table 1). UNC-8d-expressing oocytes had more positive resting membrane potentials (Table 1) following incubation in a medium containing benzamil (100 μM) in addition to amiloride (300 μM). With these findings, we replicate prior work showing that MEC-4d and UNC-8d, but not MEC-10d forms homomeric channels in oocytes (Goodman et al., 2002; Wang et al., 2013) and show that DEL-1d is not likely to form ion channels on its own.

Oocytes expressing DEGT-1d generated small currents at −85 mV and these currents reversed polarity at significantly more positive potentials than those recorded from uninjected cells and those expressing MEC-10d and DEL-1d. The reversal potential of DEGT-1d currents was also significantly more negative than those recorded from cells expressing MEC-4d and UNC-8d (Fig. 1, B-D, I and Table 1). Together, these findings suggest that DEGT-1d is sufficient to generate an ion channel whose ion permeability appears to differ from channels formed by MEC-4d and UNC-8d. These properties are not conferred by MEC-2, since DEGT-1 currents were indifferent to its presence (Table 1). Similarly, they are not conferred by MEC-2 since oocytes expressing MEC-2 along were indistinguishable from uninjected oocytes [Fig. S1 A, Table 1, and (Goodman et al., 2002)].

To verify that DEGT-1d is able to form homomeric channels, we sought recording conditions in which the current could be potentiated or blocked. As other members of this superfamily form acid-sensitive ion channels, we tested the effect of increasing the extracellular proton concentration on currents carried by DEGT-1d, MEC4-d, UNC-8d switching from solutions from pH 8.4 and 6.4. This maneuver decreased current carried by DEGT-1d and, to a lesser degree, MEC-4d, leading to a positive *I*_ΔpH_ (Fig. 1 J, top). By contrast, acidification appeared to potentiate UNC-8d currents (Fig. 1 J, top). To determine whether these effects differed from that found in control oocytes, we used an estimation approach to compute the distribution of Δ*I*_ΔpH_ (effect size and 95% confidence intervals) for each channel type (Fig. 1 J, bottom). Collectively, these findings indicate that DEGT-1d forms a homomeric channel with properties that differ from most other DEG/ENaC/ASIC channels and suggest that alkalization could enhance and acidification could suppress DEGT-1-dependent currents *in vivo*.

**Table 1:**
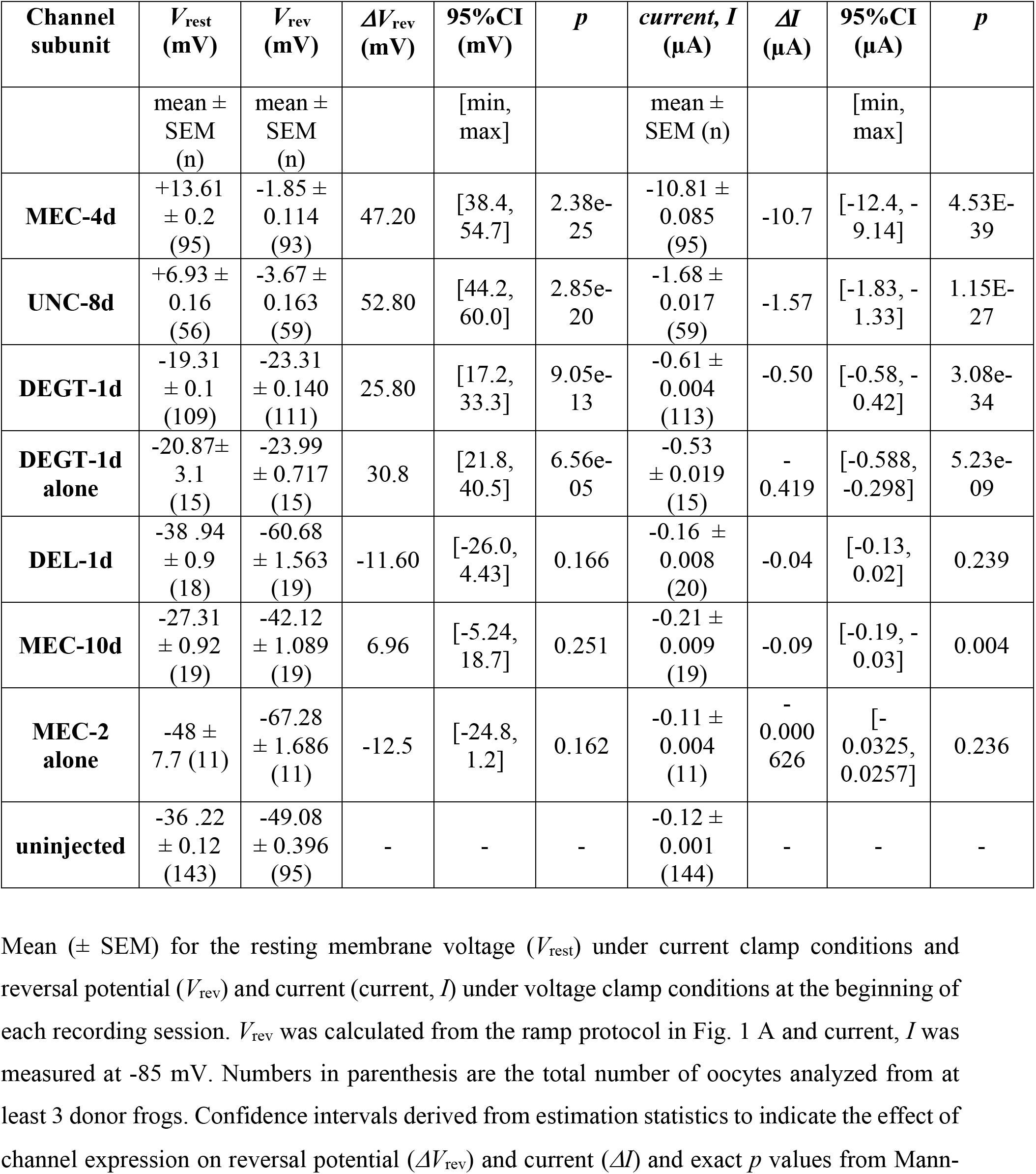
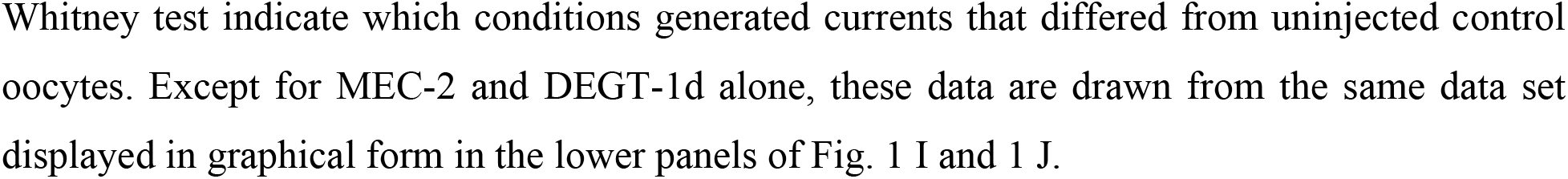
Resting membrane potential and properties of membrane current in oocytes expressing DEG/ENaC/ASIC subunits.

| Channel subunit | $V_{rest}$ (mV) | $V_{rev}$ (mV) | $\Delta V_{rev}$ (mV) | 95%CI (mV) | $p$ | current, $I$ (μA) | $\Delta I$ (μA) | 95%CI (μA) | $p$ |
| --- | --- | --- | --- | --- | --- | --- | --- | --- | --- |
| | mean ± SEM (n) | mean ± SEM (n) | | [min, max] | | mean ± SEM (n) | | [min, max] | |
| **MEC-4d** | +13.61 ± 0.2 (95) | -1.85 ± 0.114 (93) | 47.20 | [38.4, 54.7] | 2.38e-25 | -10.81 ± 0.085 (95) | -10.7 | [-12.4, -9.14] | 4.53E-39 |
| **UNC-8d** | +6.93 ± 0.16 (56) | -3.67 ± 0.163 (59) | 52.80 | [44.2, 60.0] | 2.85e-20 | -1.68 ± 0.017 (59) | -1.57 | [-1.83, -1.33] | 1.15E-27 |
| **DEGT-1d** | -19.31 ± 0.1 (109) | -23.31 ± 0.140 (111) | 25.80 | [17.2, 33.3] | 9.05e-13 | -0.61 ± 0.004 (113) | -0.50 | [-0.58, -0.42] | 3.08e-34 |
| **DEGT-1d alone** | -20.87 ± 3.1 (15) | -23.99 ± 0.717 (15) | 30.8 | [21.8, 40.5] | 6.56e-05 | -0.53 ± 0.019 (15) | -0.419 | [-0.588, -0.298] | 5.23e-09 |
| **DEL-1d** | -38.94 ± 0.9 (18) | -60.68 ± 1.563 (19) | -11.60 | [-26.0, 4.43] | 0.166 | -0.16 ± 0.008 (20) | -0.04 | [-0.13, 0.02] | 0.239 |
| **MEC-10d** | -27.31 ± 0.92 (19) | -42.12 ± 1.089 (19) | 6.96 | [-5.24, 18.7] | 0.251 | -0.21 ± 0.009 (19) | -0.09 | [-0.19, -0.03] | 0.004 |
| **MEC-2 alone** | -48 ± 7.7 (11) | -67.28 ± 1.686 (11) | -12.5 | [-24.8, 1.2] | 0.162 | -0.11 ± 0.004 (11) | -0.000626 | [-0.0325, 0.0257] | 0.236 |
| **uninjected** | -36.22 ± 0.12 (143) | -49.08 ± 0.396 (95) | - | - | - | -0.12 ± 0.001 (144) | - | - | - |

**Table 2:** Effect of amiloride analogs and NSAIDs on DEGT-1d currents relative to uninjected oocytes: estimation statistics supporting Fig. 3 and Fig. 5.

| DEGT-1d | $\Delta I_{\text{drug}}$ ($\mu\text{A}$) | 95% CI ($\mu\text{A}$) | *p* |
| --- | --- | --- | --- |
| *amiloride analogs* |  |  |  |
| amiloride | 0.0033 (20) | [-0.02, 0.025] | 0.36 |
| benzamil | 0.0073 (18) | [-0.009, 0.035] | 0.61 |
| EIPA | 0.0073 (13) | [-0.008, 0.028] | 0.43 |
| phenamil | -0.0043 (13) | [-0.024, 0.016] | 0.32 |
| benzamidine | 0.0041(13) | [-0.012, 0.026] | 0.77 |
| *NSAIDs* |  |  |  |
| ibuprofen | 0.02 (18) | [0.0068, 0.036] | 0.001 |
| flurbiprofen | 0.15 (5) | [0.084, 0.22] | 0.003 |
| diclofenac | -0.037 (5) | [-0.075, -0.015] | 0.003 |
| aspirin | 0.037 (5) | [0.0075, 0.064] | 0.08 |
| salicylic acid | 0.073 (4) | [0.042, 0.094] | 0.006 |

**Table 3:** Effect of amiloride analogs and NSAIDs on UNC-8d currents relative to uninjected oocytes: estimations statistics supporting Fig. 3 and Fig. 5.

| UNC-8d | $\Delta I_{\text{drug}}$ (uA) | 95% CI ($\mu\text{A}$) | *p* |
| --- | --- | --- | --- |
| *amiloride analogs* |  |  |  |
| amiloride | 0.22 (5) | [0.026, 0.29] | 0.03 |
| benzamil | 0.87 (18) | [0.64, 1.13] | 2.08e-09 |
| EIPA | 0.61 (9) | [0.45, 0.83] | 4.92e-06 |
| phenamil | 0.18 (10) | [0.12, 0.26] | 2.86e-06 |
| benzamidine | 0.32 (9) | [0.17, 0.58] | 1.29e-05 |
| *NSAIDs* |  |  |  |
| ibuprofen | 0.026 (15) | [0.000518, 0.0492] | 0.024 |
| flurbiprofen | 0.22 (10) | [0.123, 0.382] | 0.00033 |
| diclofenac | 0.24 (13) | [0.156, 0.352] | 5.25e-06 |
| aspirin | -0.071 (9) | [-0.162, 0.00864] | 0.39 |
| salicylic acid | -0.062 (10) | [-0.197, 0.00931] | 0.16 |

**Table 4:**
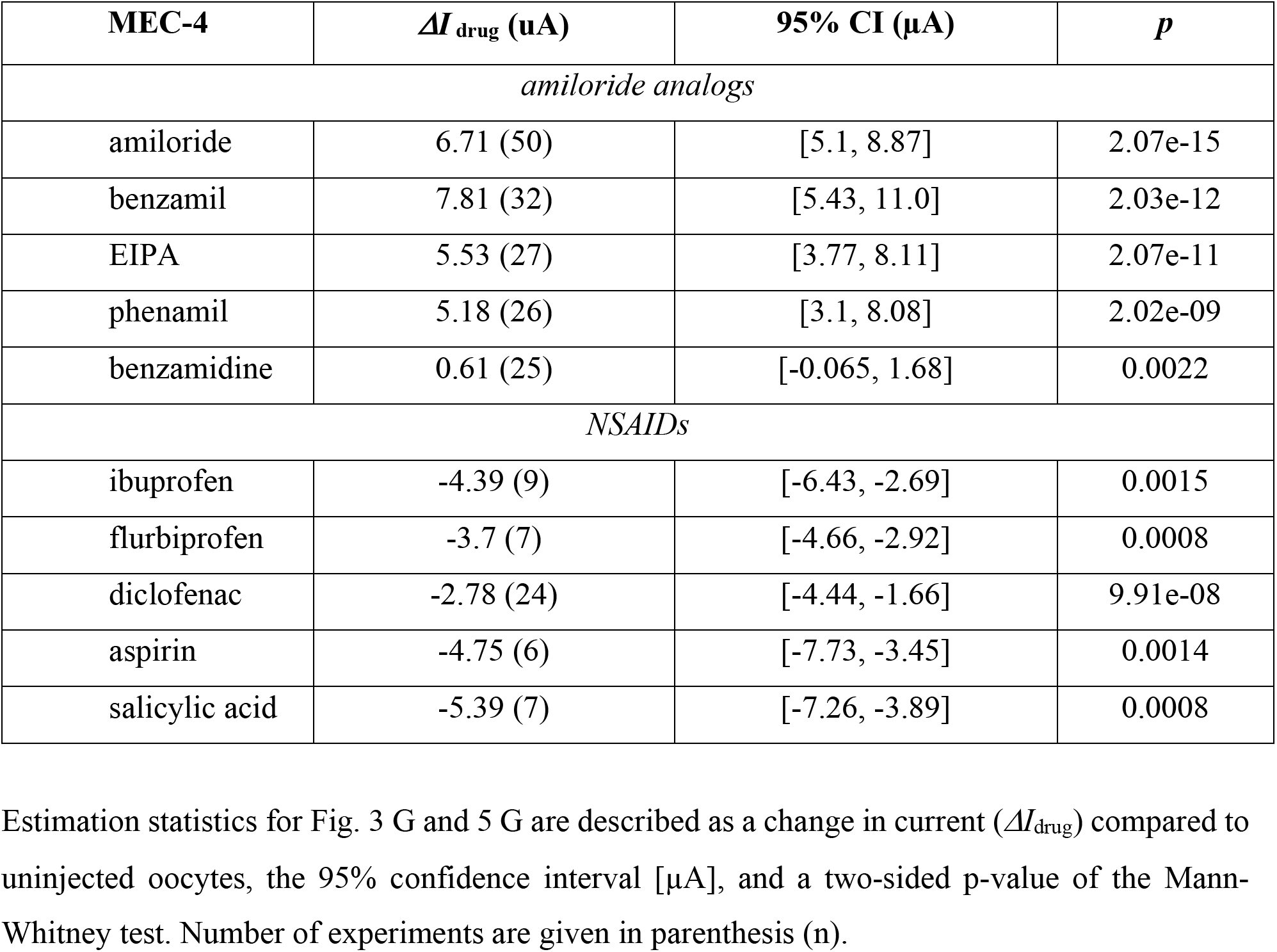
Effect of amiloride analogs and NSAIDs on MEC-4d currents relative to uninjected oocytes: estimations statistics supporting Fig. 3 and Fig. 5.

| MEC-4 | $\Delta I_{\text{drug}}$ (uA) | 95% CI ($\mu\text{A}$) | *p* |
| --- | --- | --- | --- |
| *amiloride analogs* |  |  |  |
| amiloride | 6.71 (50) | [5.1, 8.87] | 2.07e-15 |
| benzamil | 7.81 (32) | [5.43, 11.0] | 2.03e-12 |
| EIPA | 5.53 (27) | [3.77, 8.11] | 2.07e-11 |
| phenamil | 5.18 (26) | [3.1, 8.08] | 2.02e-09 |
| benzamidine | 0.61 (25) | [-0.065, 1.68] | 0.0022 |
| *NSAIDs* |  |  |  |
| ibuprofen | -4.39 (9) | [-6.43, -2.69] | 0.0015 |
| flurbiprofen | -3.7 (7) | [-4.66, -2.92] | 0.0008 |
| diclofenac | -2.78 (24) | [-4.44, -1.66] | 9.91e-08 |
| aspirin | -4.75 (6) | [-7.73, -3.45] | 0.0014 |
| salicylic acid | -5.39 (7) | [-7.26, -3.89] | 0.0008 |

### Unlike MEC-4d and UNC-8d, DEGT-1d is insensitive to amiloride analogs

The DEG/ENaC/ASIC ion channel family is also known as the amiloride-sensitive ion channel (ASC) family (Goodman and Schwarz, 2003; Hanukoglu and Hanukoglu, 2016; Kellenberger and Schild, 2002), suggesting that channels formed by these proteins are sensitive to the diuretic amiloride and its derivatives. Indeed, both MEC-4d and UNC-8d are known to be blocked by amiloride (Goodman et al., 2002; Wang et al., 2013). MEC-4d has a micromolar affinity for amiloride and UNC-8d is ~8-fold less sensitive to amiloride (Brown et al., 2007; Goodman et al., 2002; Wang et al., 2013). To learn more about amiloride block as a shared, but potentially variable property of DEG/ENaC/ASIC channels, we tested DEGT-1d, UNC-8d, and MEC-4d for sensitivity to amiloride (Amil) and four analogs: benzamil (Bmil), 5-(N-Ethyl-N-isopropyl)amiloride (EIPA), phenamil (Phen), and benzamidine (Bzd) (Fig. 2 and 3). These analogs were developed in an effort to generate specificity for ENaCs and Na^+^/H^+^ antiporters, both of which play critical roles in the mammalian kidney and are inhibited by amiloride (Frelin et al., 1987).

**Figure 2:**
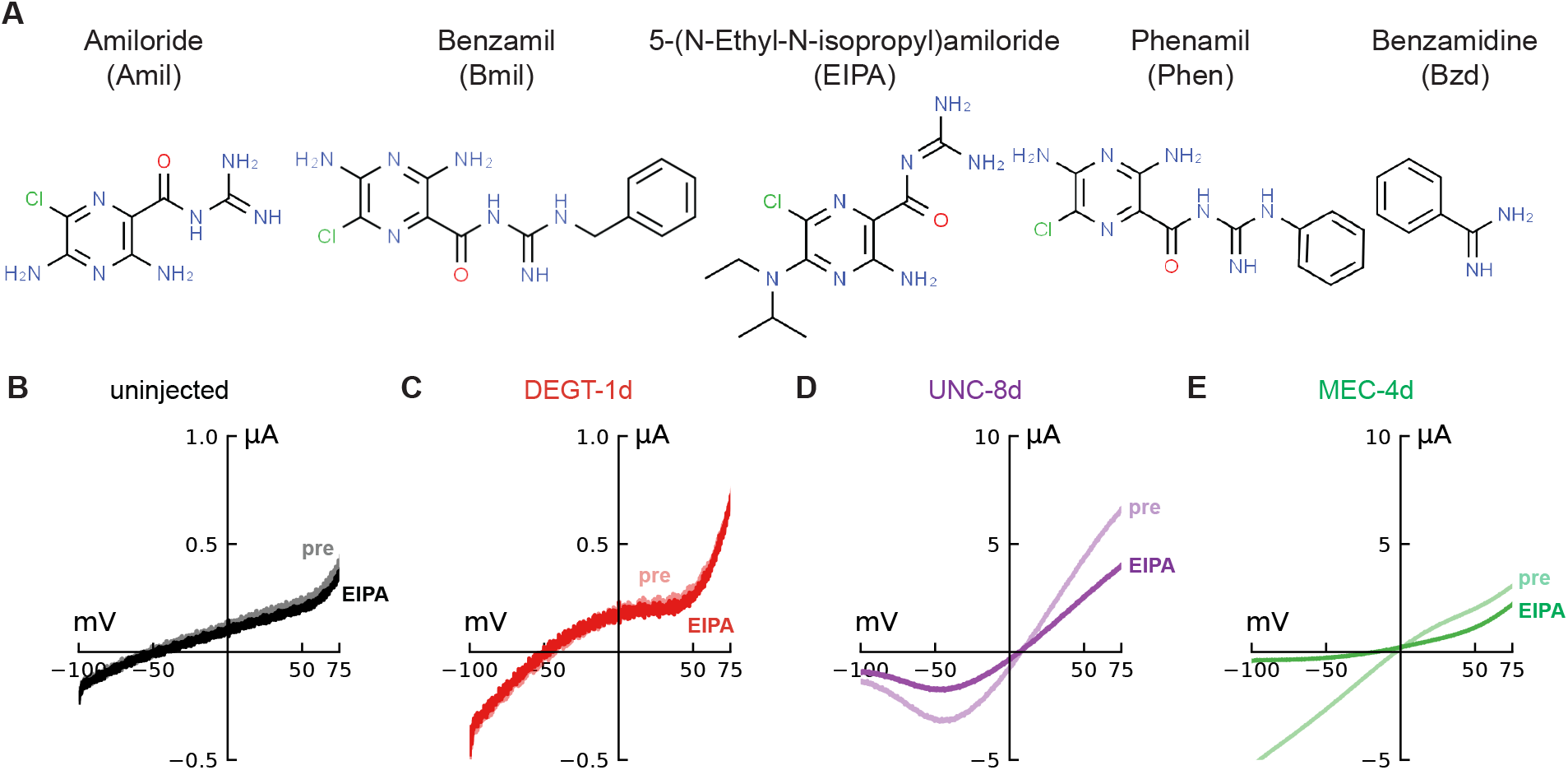
Structures of amiloride analogs and representative current-voltage curves in the absence (pre) and presence of 5-(N-Ethyl-N-isopropyl) amiloride (EIPA). **(A)** Structures of amiloride analogs used in this study. **(B-E)** Current-voltage curves of uninjected oocytes (**B**) and oocytes co-expressing MEC-2 and DEGT-1d (**C**), UNC-8d (**D**) or MEC-4d (**E**) in the absence (lighter color, pre) and presence (darker color, EIPA) of 30 μM EIPA.

**Figure 3:**
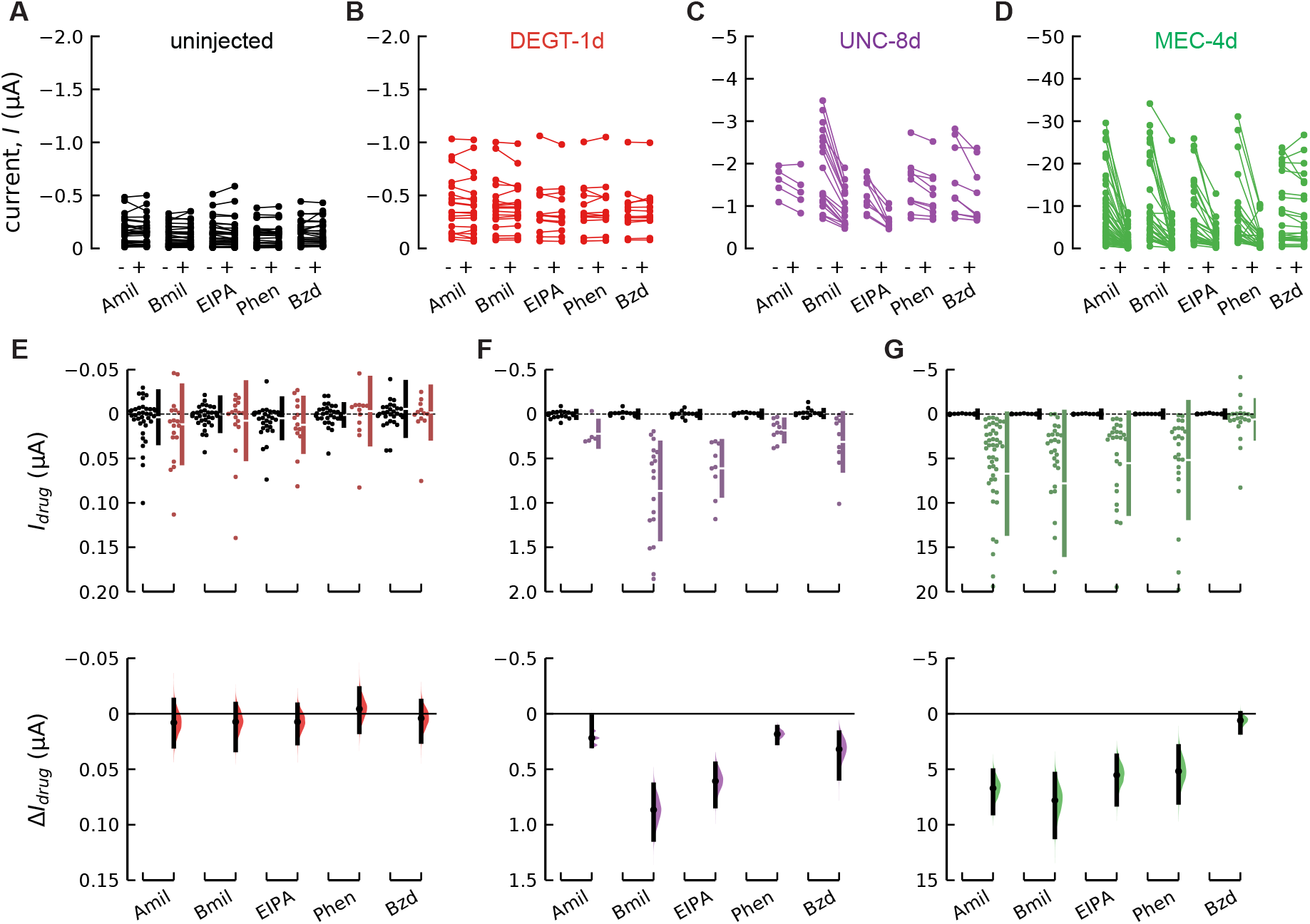
Unlike MEC-4d and UNC-8d, DEGT-1d currents are insensitive to amiloride analogs. **(A-D)** Paired dots show the current at −85 mV in uninjected oocytes **(A)** and in oocytes expressing DEGT-1d **(B)**, UNC-8d **(C)** or MEC-4d **(D,)** before and after treatment with 30 μM amiloride (Amil), benzamil (Bmil), EIPA, phenamil (Phen) or benzamidine (Bzd). **(E-G)** Drug-sensitive current (*I*_drug_) (top) at −85 mV in oocytes expressing DEGT-1d **(E)**, UNC-8d **(F)** or MEC-4d **(G)** (in color) compared to uninjected oocytes (black). Estimation plots (bottom) showing the effect of each drug (*ΔI*_drug_) on DEGT-1d **(E)**, UNC-8d **(F)** or MEC-4d **(G)** relative to the drug effect on uninjected oocytes. Mean values, 95% confidence intervals, and statistical analyses related to panels **E-G** are in Table 2 – 4.

For simplicity, we exposed oocytes expressing DEGT-1d, UNC-8d, and MEC-4d to a single concentration (30 μM) of each amiloride analog. Figure 2, B-E show representative current-voltage curves recorded in the presence and absence of one amiloride analogue, EIPA, in uninjected (control) oocytes and in oocytes expressing each channel. EIPA had no detectable effect on currents in control oocytes or in those expressing DEGT-1d (Fig. 2, A and B). By contrast, EIPA inhibited both UNC-8d and MEC-4d current (Fig. 2, D and E). During each recording, we also measured current amplitude at −85 mV (see voltage protocol Fig. 1 A, top). Figure 3, A-D shows the average current measured in the presence and absence of each analog for individual control, DEGT-1d, UNC-8d, and MEC-4d oocytes. Next, we determined the drug-sensitive difference current, *I*_drug_, for each channel isoform and for control oocytes (Fig. 3, E-G, top). Finally, we adopted an estimation statistics approach (Ho et al., 2019) to determine the size of the drug effect, Δ*I*_drug_ on DEGT-1d, UNC-8d, and MEC-4d relative to control (Fig. 3, E-G, bottom). In this representation, negligible effects result in a distribution centered near 0 and inhibitory effects shift the distribution to more positive values.

Unlike MEC-4d, UNC-8d and most other DEG/ENaC/ASIC channels, DEGT-1d was insensitive to all amiloride analogs we tested at our reference concentration of 30 μM (Fig. 3, B and E). To differentiate between a reduced affinity and a lack of sensitivity, we tested concentrations up to 300 μM for amiloride. At this concentration, amiloride reduced current in DEGT-1 expressing oocytes by 60 nA, on average, relative to its effect on control oocytes (Fig. S1 A).

UNC-8d is inhibited not only by amiloride and benzamil (Wang et al., 2013, Miller-Flemming et al., 2016), but also by EIPA, phenamil and benzamidine (Fig. 3, C and F). Amiloride analogs block UNC-8d currents with different degrees of potency. In ascending order of potency, UNC-8d was blocked by amiloride, phenamil, benzamidine, EIPA and benzamil. The reported *IC*_50_ values for UNC-8d channels for amiloride at −100 mV are 7.8 μM in divalent-containing and 106 μM in divalent free solution (Wang et al., 2013). The reported *IC*_50_ values for UNC-8d channels for benzamil at −100 mV are 47 μM in divalent-containing and 119 μM in divalent free solution (Miller-Flemming et al., 2016). In this study, we determined an *IC*_50_ for benzamil in control saline (divalent solution) of 14.8 ± 1.6 μM (n =4) at −60 mV (15.1 μM at −100 mV) (Fig. S2, A-C). In contrast to amiloride inhibition, the apparent affinity to benzamil was indistinguishable at voltages between −100 and −20 mV, indicating that the block through benzamil is not voltage-dependent.

Similar to UNC-8d, MEC-4d is inhibited by amiloride, benzamil and benzamidine (Brown et al., 2007) and by EIPA and phenamil (Fig. 3, D and G). However, the order of potency differs from UNC-8d. In ascending order: MEC-4d was blocked by benzamidine < amiloride ≈ EIPA ≈ phenamil < benzamil. The similar potency of amiloride and EIPA was unexpected and we analyzed this drug further by collecting full dose-response curves for EIPA inhibition of MEC-4d (Fig. S2, D-F). The half-blocking dose or *IC*_50_ for EIPA was 3.06 ± 0.6 μM (n = 11) at −60 mV (Fig. S2, E-F), which is indistinguishable from the *IC*_50_ for amiloride (2.35 ± 0.39 μM (n = 7) at −60 mV (Brown et al., 2007). It has been shown that the block through amiloride in ENaC, MEC-4d and UNC-8d depends on the transmembrane potential difference such that hyperpolarization of the membrane increases channel block (Kellenberger & Schild 2002, Brown 2007, Wang 2013). By comparison, the apparent affinity to EIPA was insensitive to voltage (Fig. S2 F). Collectively, these results show that MEC-4d is blocked by many amiloride analogs, in order of potency: benzamidine (196 μM) < EIPA (3.06 μM, this study) ≈ amiloride (2.35 μM) ≈ phenamil (TBD) < benzamil (0.83 μM) (Brown, 2007).

If all of these drugs were to function as open channel blockers like amiloride (Brown et al., 2007; Waldmann et al., 1995), then the dramatic difference in their potency among DEGT-1d, UNC-8d and MEC-4d channels implies that these channels differ in the molecular pathways by which drugs access a common binding site in the pore or that present distinct binding sites. We favor the former idea for two reasons. First, the conserved second transmembrane domain of DEG/ENaC/ASIC proteins has long been thought to line the ion conduction pore and to present a conserved amiloride binding site represented (Kellenberger et al., 1999; Snyder et al., 1999). Second, access to this site in the pore would be influenced by differences in the open-state conformation of the extracellular domain and this is the region that is most divergent among the channels we studied.

### Some NSAIDs block DEGT-1d and UNC-8d, but all potentiate MEC-4d

Because nonsteroidal anti-inflammatory drugs (NSAIDs) have been reported to block ASIC channels with *IC*_50_ values in the high micromolar range (90-350 μM) (Lingueglia and Lazdunski, 2013; Voilley, 2004), we examined sensitivity to five NSAID drugs (Fig. 4 A) as an additional window into shared, but variable properties of DEG/ENaC/ASIC channels. Current-voltage curves of uninjected (control), DEGT-1d, UNC-8d, and MEC-4d currents (Fig. 4, B-E) show that ibuprofen has little, if any effect on control currents, modestly inhibits DEGT-1d and UNC-8d, and seems to potentiate MEC-4d. None of the NSAIDS tested affected currents in control oocytes at 30 μM (Fig. 5 A), providing a simple background for assessing their effect on DEGT-1d, UNC-8d and MEC-4d.

**Figure 4:**
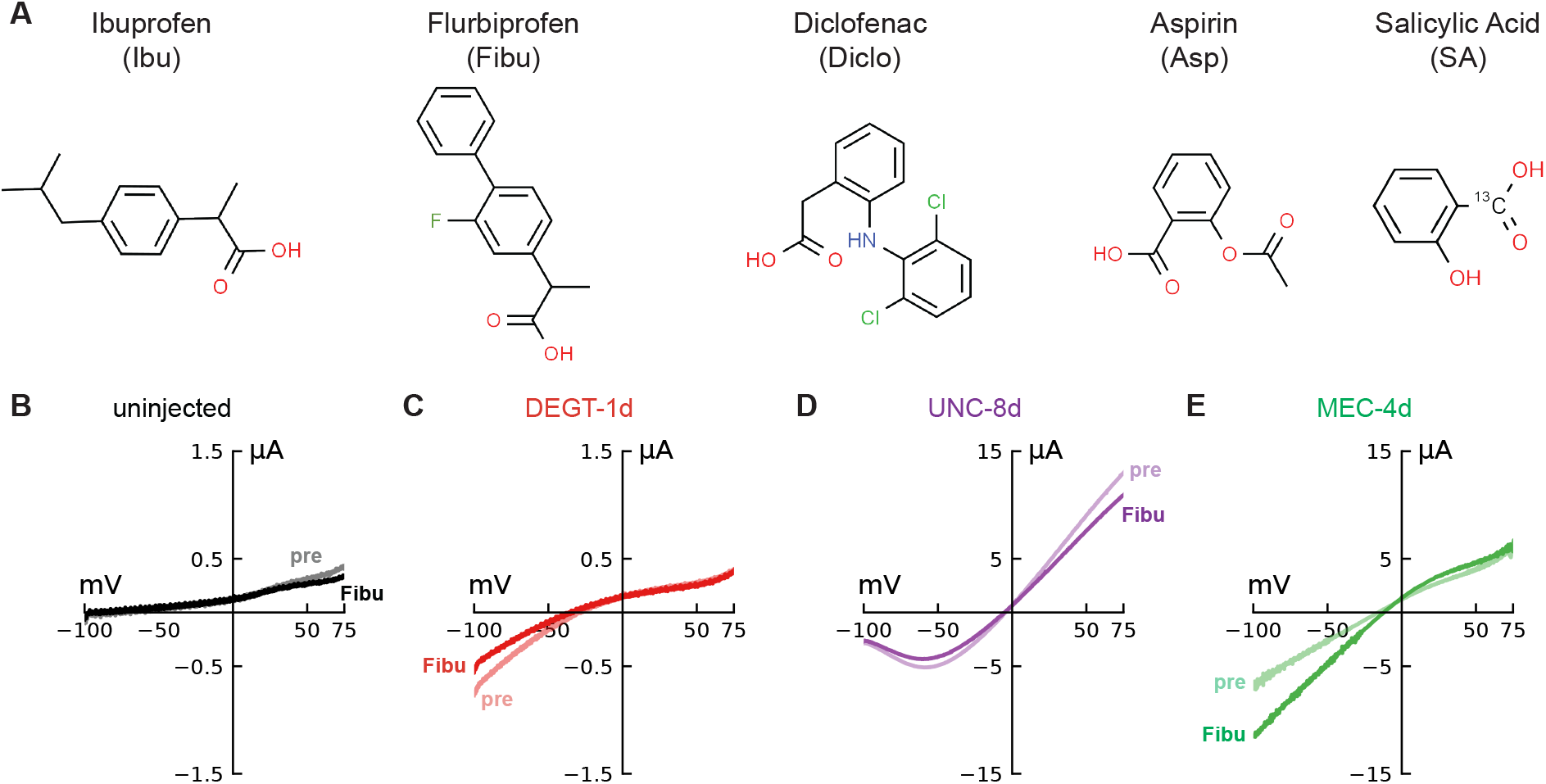
Structures of NSAIDs and representative current-voltage curves in the absence (pre) and presense of fluoribuprofen (Fibu). **(A)** Chemical structures of NSAIDs used in this study. **(B-E)** Current-voltage curves of uninjected oocytes (**B**) and oocytes expressing DEGT-1d (**C**), UNC-8d (**D**) or MEC-4d (**E**) in the absence (lighter color) and presence (darker color) of 30 μM Flurbiprofen (Fibu).

**Figure 5:**
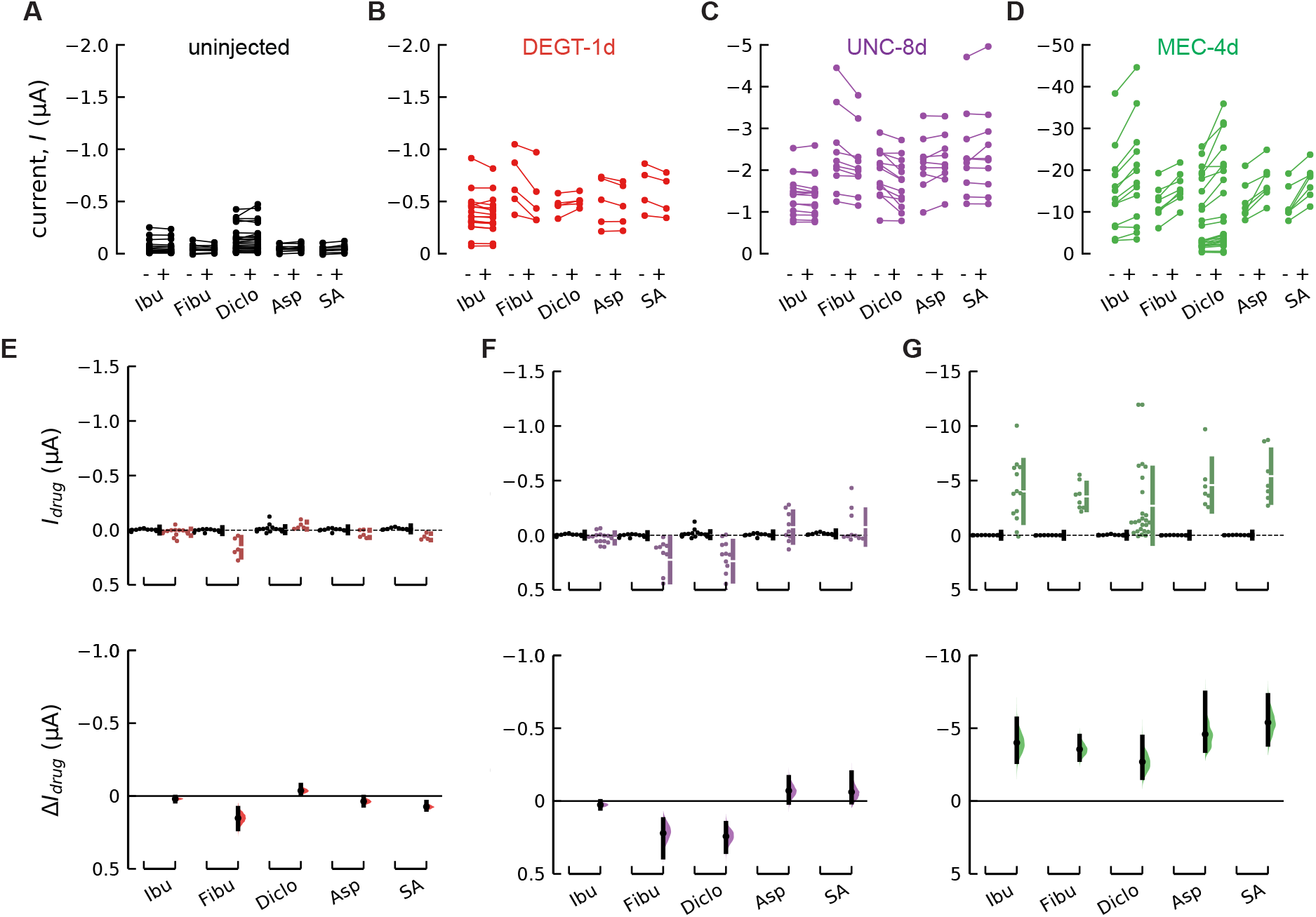
Nonsteroidal anti-inflammatory drugs (NSAIDS) potentiate MEC-4d current and inhibit or are ineffective on DEGT-1d and UNC-8d. **(A-D)** Paired dots show the current at 85 mV in uninjected oocytes **(A)** and in oocytes co-expressing MEC-2 and DEGT-1d **(B)**, UNC-8d **(C)** or MEC-4d **(D)** before and after treatment with 30 μM ibuprofen (Ibu), Flurbiprofen (Fibu), Diclofenac (Diclo), Aspirin (Asp), salicylic acid (SA). **(E-G)** Drug-sensitive current (*I*_drug_) (top) at −85 mV in oocytes expressing DEGT-1d **(E)**, UNC-8d **(F)** or MEC-4d **(G)** (in color) compared to uninjected oocytes (black). Estimation plots (bottom) showing the effect of each drug (*ΔI*_drug_) on DEGT-1d **(E)**, UNC-8d **(F)** or MEC-4d **(G)** relative to the drug effect on uninjected oocytes. Mean values, 95% confidence intervals, and statistical analyses related to panels **E-G** are in Table 2–4.

Similar to our strategy for analyzing the effect of amiloride analogs, we measured current (at −85 mV) in the absence and presence of each NSAID and plotted these paired values for control, DEGT-1d, UNC-8d, and MEC-4d currents (Fig. 5, A-D). Next, we used these data to determine the drug-sensitive current, *I*_drug_, in control and channel-expressing oocytes (Fig. 5, E-G, top) and estimation statistics to determine if the effects exceeded those expected for control oocytes, Δ*I*_drug_ (Fig. 5, E-G, bottom). Collectively, this analysis indicates that 30 μM flurbiprofen (Fibu) and salicylic acid (SA) partially inhibit DEGT-1(Fig. 5, B and E) and that 30 μM Fibu and diclofenac partially inhibits UNC-8d (Fig. 5, C and F). Higher concentrations of ibuprofen also blocked DEGT-1d currents (Fig. S1 B). Surprisingly, all five NSAIDs potentiated MEC-4d currents (Fig. 5, D and G). These findings demonstrate that NSAIDs can function both as antagonists or agonists of DEG/ENaC/ASIC channels, depending on the specific channel target.

Next, we applied two NSAIDs (ibuprofen and aspirin) to cells expressing MEC-4d channels and analyzed the dose-response relationship as a function of membrane voltage (Fig. 6, A-C). To improve the sensitivity of these measurements, we reduced the baseline currents by injecting 1.6-fold less cRNA encoding MEC-4d for these experiments. Figure 6A shows MEC-4d current evoked by a family of voltage steps in the absence (left) and presence (right) of aspirin. The mean *EC*_50_ (± SEM) for ibuprofen and aspirin at −100 mV are 34.6 ± 0.9 μM (n = 12) and 79.9 ± 3.7 μM (n = 9), respectively (Fig. 6 B). Neither drug showed evidence of voltage-dependence (Fig. 6 C), suggesting that the binding site for these drugs lies outside the pore region. Ibuprofen is an enantiomer containing two chiral molecules and the S-isoform is the preferred ligand for its primary targets, the cyclo-oxygenase enzymes, COX-1 and COX-2 (Orlando et al., 2015; Selinsky et al., 2001). In contrast, MEC-4d potentiation is equally sensitive to both ibuprofen enantiomers (Fig. 6 D) and is less sensitive to ibuprofen than COX-1 and COX-2 (Blobaum and Marnett, 2007). Collectively, these findings suggest that the binding sites for NSAIDs differ in DEG/ENaC/ASIC channels and cyclo-oxygenases.

**Figure 6.**
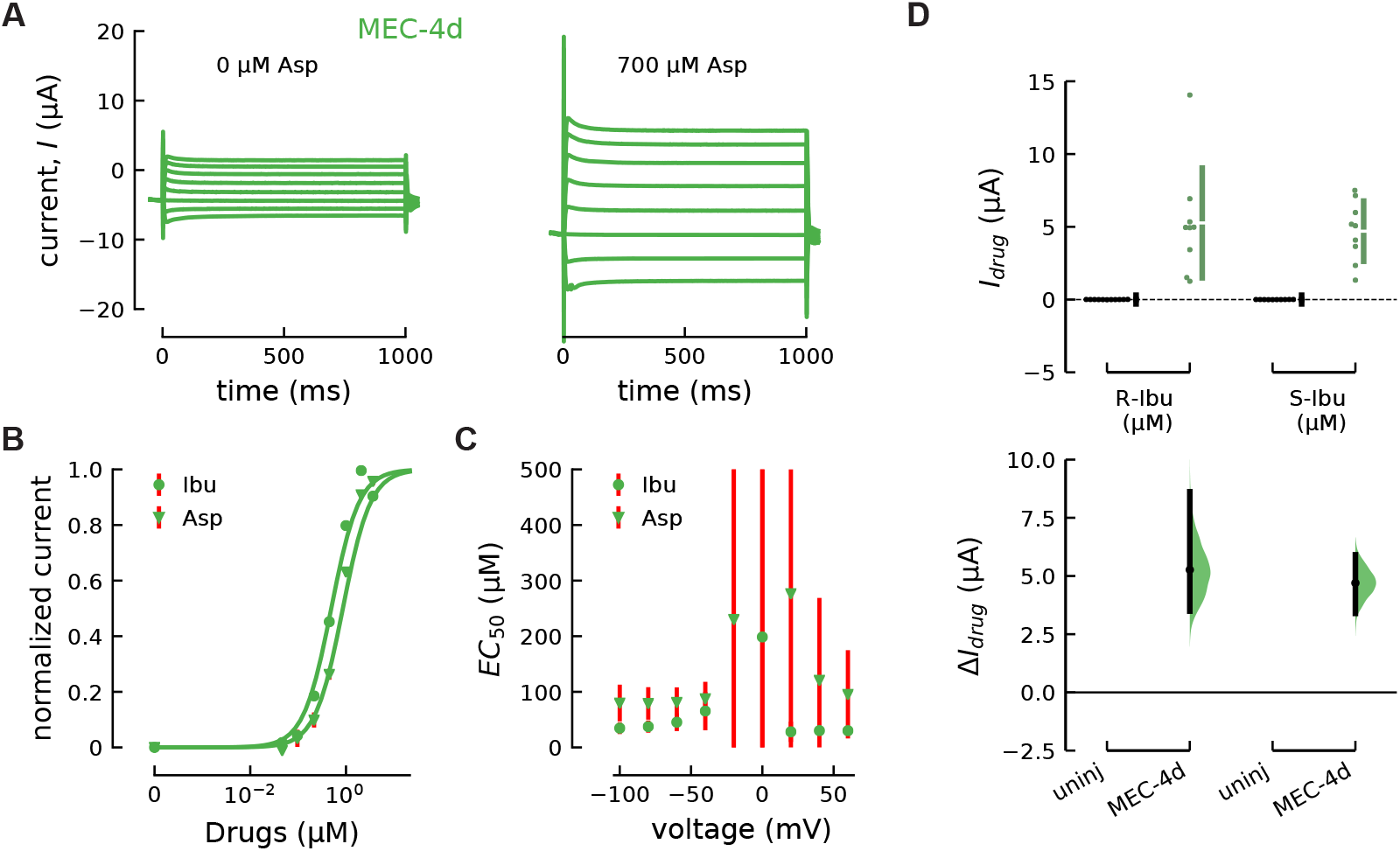
Sensitivity of MEC-4d channels to NSAIDs Ibuprofen and Aspirin. **(A)** MEC-4d current in the absence (left) and presence of aspirin (Asp, 700 μM, right). **(B)** Ibuprofen (Ibu, circles) and aspirin (Asp, triangles) dose-responses relationship at −100 mV (normalized to *I*_max_ and baseline current). Points are mean ± sem (Ibu, n=12; Asp, n = 9) and error bars to smaller than the points in most cases. **(C)***EC*_50_ values for different voltages for Ibu (circle) and Asp (triangle). **(D)** Drug-sensitive MEC-4d current (*I*_drug_) (top) and estimation plots (bottom) in the presence and absence of ibuprofen isomers (R-Ibu) and (S-Ibu) applied at 30 μM compared to uninjected oocytes (black) (***Δ****I*_drug_). Estimation plots in panel D (bottom) illustrate the 95% confidence interval [in μA], and the two-sided p-value of the Mann-Whitney test. The difference for MEC-4d for 30 μM R-Ibu is 5.27 μA [3.49, 8.62], p = 0.000197 and for 30 μM S-Ibu is 4.7 μA [3.39, 5.91], p = 0.00028.

**Figure 7:**
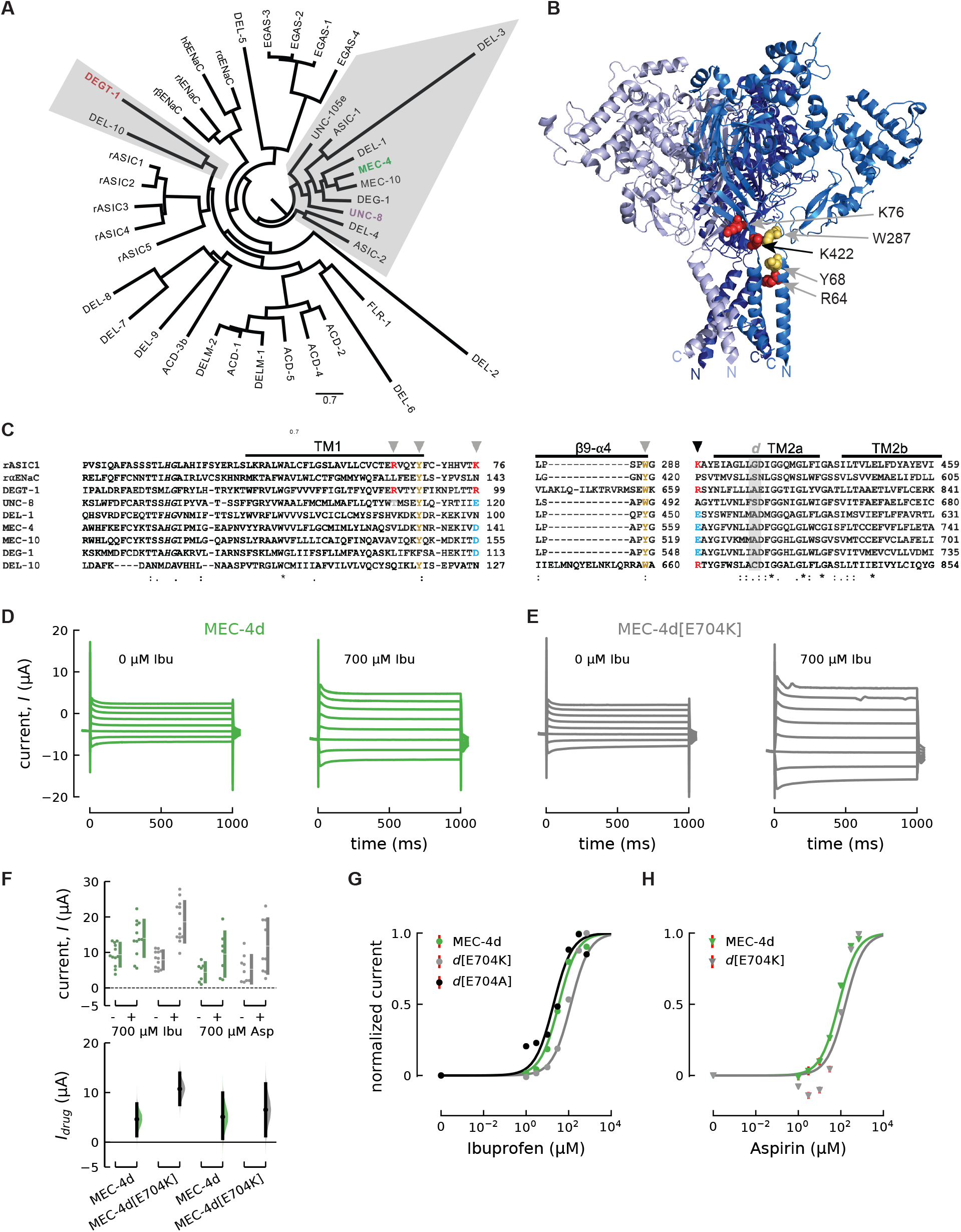
Amino acid in the wrist close to TM2 in MEC-4d changes sensitivity to ibuprofen and aspirin. **(A)** Phylogenetic tree of *C. elegans* DEG/ENaC/ASIC subunits and mammalian ENaC and ASIC subunits. Accession numbers are given in Material and Methods. **(B)** Ribbon diagram of trimeric cASIC1a (PDB 4NTW) rendered in PyMOL. Residues shown in space filling mode are those linked to ibuprofen binding in rASIC1a (Lynagh et al., 2017). **(C)** Amino acid alignment of rASIC1, rαENaC, DEGT-1, UNC-8, MEC-4, DEL-1, MEC-10, DEG-1 and DEL-10 made with Clustal Omega. (left) N-terminal domain and transmembrane domain 1 (TM1); (right) ß9-α4 and transmembrane domain 2 (TM2). Secondary structure motifs numbered as in (Jasti et al., 2007). rASIC1a[K422] (black arrowhead) implicated in Ibu sensitivity. The grey *d* indicates the site that mutates to cause degeneration in *C. elegans*. Amino acids implicated in rASIC1 ibuprofen responses are highlighted as follows: red (positively charged), blue (negatively charged) and yellow (hydrophobic). **(D)** MEC-4d current traces and **(E)** MEC-4d[E704K] current in the absence (left) and presence of 700 μM (right) ibuprofen (Ibu). **(F)** Current amplitude (current, *I*) (top) at −85 mV in the presence and absence of ibuprofen (Ibu) and Aspirin (Asp) for the MEC-4d isoform (green) and the MEC-4d[E704K] isoform (right). Estimation plots (bottom) showing the drug-induced change in current (*I*_*drug*_) for MEC-4d (green) or MEC-4d[E704K] (grey) in the presence of 700 μM Ibu or aspirin (Asp) compared to the absence of the drugs. Estimation plots show the 95% confidence interval [in μA], and the two-sided p-value of the Mann-Whitney test. The effect of 700 μM Ibu is 4.63 μA [1.29, 7.73], p = 0.0102 (n = 12) and 10.7 μA [7.58, 13.9], p = 5.09e-05 (n = 13) for MEC-4d and and MEC-4d[E704K], respectively. The effect of 700 μM Asp is 5.11 μA [0.74, 9.92], p = 0.073 (n = 7) and 6.53 μA [1.28, 11.8], p = 0.0521 (n = 9) for MEC-4d and MEC-4d[E704K], respectively. **(G)** Ibuprofen dose-responses relationships for MEC-4d (green), MEC-4d[E704K] (grey), and MEC-4d[E704A] (black) currents at −100 mV (normalized to *I*_max_ and baseline current). **(H)** Aspirin dose-responses relationships for MEC-4d (green) and MEC-4d-E704K (grey) currents at −100 mV (normalized to *I*_max_ and baseline current).

MEC-4d is not the only member of the large DEG/ENaC/ASIC ion channel family (Fig. 7 A) affected by NSAIDs. rASIC1a is inhibited by ibuprofen and this allosteric effect depends on three positively charged and two hydrophobic residues near TM1 and TM2 (Fig. 7, B and C) (Lynagh et al., 2017). The putative binding site for ibuprofen is proposed to include these five residues. To learn more, we aligned and compared the sequences of seven *C. elegans* DEG/ENaC/ASIC channels with rASIC1a (Fig. 7 C). We found that MEC-4 differs from rASIC1a at the three of the five residues linked to inhibition by ibuprofen (Fig. 7 C, arrowheads). Here, we focused on K422 in rASIC1a and E704, the glutamate at the homologous position in MEC-4 (Fig. 7 C, black arrowhead). Comparing the effect of ibuprofen on MEC-4d and MEC-4d[E704K], we found that ibuprofen potentiated both isoforms (Fig. 7, D and E). We quantified this effect by collecting dose-response curves for ibuprofen and aspirin (Fig. 7 G and H). Sensitivity to ibuprofen was modestly increased in MEC-4d[E704K], but unaffected in MEC-4d[E704A]: the mean *EC*_50_ (± SEM) to ibuprofen for MEC-4d[E704K] and MEC-4d[E704A] at −100 mV are 135 ± 40 μM (n = 13) and 19.5 ± 8.84 μM (n = 14), respectively (Fig. 7 G). A similar shift was observed for potentiation for MEC-4d[E704K] by aspirin with a mean EC_50_ (± SEM) of 179 ± 56 μM (n = 9) (Fig. 7 H). This finding differs from allosteric inhibition of rASIC1a, which is significantly impaired by introducing alanine into this position (Lynagh et al., 2017). In an effort to discover the domains responsible for these differences between rASIC1a and MEC-4d, we designed constructs encoding chimeras of these two channels. These chimeras did not generate any detectable current, however, even when co-expressed with MEC-2. The ability of ibuprofen to potentiate MEC-4d and to inhibit rASIC1a could reflect the existence of distinct binding sites in the two channel isoforms or a common, conserved binding site and distinct energetic coupling between ibuprofen binding and channel gating. Based on the similar effects of mutagenesis on apparent ibuprofen affinity, we favor the simpler model in which NSAIDs share a similar binding site in MEC-4d and rASIC1a. Future studies to directly determine the ibuprofen binding sites will be required to differentiate between these classes of models, however.

## Discussion

### Sensitivity to amiloride and its analogs

Developed in 1967 to treat hypertension, amiloride is listed an essential medicine by the world health organization (World Health Organization, 2019). Many amiloride derivatives have been developed and we leveraged this collection to evaluate the sensitivity of DEGT-1d, UNC-8d and MEC-4d to a panel of five amiloride analogs (Fig. 2 and 3). Whereas both MEC-4d and UNC-8d were inhibited by at least one amiloride analog, DEGT-1d was not obviously affected by any of the five amiloride analogs we tested. UNC-8d currents differed from MEC-4d in their sensitivity to amiloride analogs. In particular, UNC-8d currents are more sensitive to inhibition by benzamil and EIPA than to either amiloride or phenamil. Four of the five compounds inhibited currents carried by MEC-4d at a constant dose of 30 μM. [Benzamidine had little or no effect on MEC-4d at this concentration, but does block MEC-4d currents at higher doses (Brown et al., 2007).] Benzamil was the most potent inhibitor of MEC-4d currents, followed by amiloride, EIPA, and phenamil (Fig. 3 G, (Brown et al., 2007). Single-channel recordings demonstrate that amiloride functions as an open channel blocker of MEC-4 channels (Brown et al., 2007), indicating that amiloride binds within the ion conduction pore. This idea is reinforced by three-dimensional co-crystal structures of cASIC1a and amiloride revealing an amiloride molecule lodged near the external vestibule of the central pore (Baconguis et al., 2014). Together, our findings suggest that DEGT-1d, UNC-8d, and MEC-4d proteins form homomeric channels that differ in the structure of the amiloride binding site or in the accessibility of compounds to this site.

### Sensitivity to NSAIDs and their analogs

The mechanism by which non-steroidal anti-inflammatory drugs or NSAIDs generate analgesia is by inhibiting cyclo-oxygenase-1 (COX-1) and COX-2 enzymes (Day and Graham, 2013; Weissmann, 1991). These compounds also inhibit DEG/ENaC/ASIC channels (Lingueglia and Lazdunski, 2013; Lynagh et al., 2017; Voilley, 2004; Voilley et al., 2001) and P2X channels (Lynagh et al., 2017). We built on these observations and tested *C. elegans* DEG/ENaC/ASICs for sensitivity to a panel of five NSAIDs (Fig. 4 and 5). One of the five NSAIDs modestly inhibited DEGT-1d (Fig. 5 E) and UNC-8d (Fig. 5 F). By contrast with our finding and the well-characterized inhibition of ASIC1a by NSAIDs, all five compounds strongly activated MEC-4d currents (Fig. 5 G).

Ibuprofen potentiates MEC-4d currents in a dose-dependent manner (Fig. 6 B) and functions as a negative allosteric modulator of proton-gated ASIC1a currents (Lynagh et al., 2017). Based on our finding that mutating E704 decreases the apparent affinity for ibuprofen (Fig. 7 G), we propose that MEC-4d shares an ibuprofen binding site with ASIC1a. This raises the question of how ibuprofen binding might enhance MEC-4d current and suppress ASIC1a current. In ASIC1a, ibuprofen and protons elicit opposing conformational changes at the top of the pore-lining second transmembrane domain (Lynagh et al., 2017), supporting the idea that ibuprofen is a negative allosteric modulator of proton-dependent ASIC1a gating. If a similar conformational change were associated with NSAID binding to MEC-4d, then it would be uncoupled to proton binding (MEC-4d is not activated by protons) and we would infer that the motion is associated with an increase in channel gating. Future work will be needed to resolve the exact nature of the allosteric interactions between ibuprofen binding and channel gating, however. The differential response of MEC-4d and ASIC1a presents an avenue for further study.

### Concluding remarks

The DEG/ENaC/ASIC channels differ from most, if not all other classes of ion channels: they are only present in metazoan genomes (Katta et al., 2015; Liebeskind et al., 2015). Phylogenetic studies indicate that this gene superfamily has undergone expansions within certain animal lineages, including nematodes and insects (Liebeskind et al., 2015). By analyzing a subset of *C. elegans* DEG/ENaC/ASIC proteins, we extend understanding of the functional diversification of this ion channel superfamily. In particular, we show that DEGT-1 appears to lack sensitivity to amiloride and four of its derivatives. To our knowledge, this is the first member of this family to have these properties. This finding suggests that using only amiloride might well obscure the contribution of DEG/ENaC/ASIC channels to cell and tissue function. From the subset of DEG/ENaC/ASIC studied, DEGT-1 phylogenetically distant from the others (Fig. 7A). We also identified NSAIDs as potential inhibitors of DEGT-1d and UNC-8 currents and positive activators of MEC-4d currents. Thus, ibuprofen might serve as a tool to screen for the activity of other DEG/ENaC/ASICs in heterologous cells or in their native tissue. Collectively, we demonstrate that each of the proteins able to form homomeric channels in Xenopus oocytes exhibits a unique pharmacological footprint within two drug families. This property opens the door to using sensitivity to amiloride and ibuprofen to determine the composition of heterotrimeric DEG/ENaC/ASIC channels either in heterologous cells or in their native tissues.

## Acknowledgements

We thank Z. Liao for excellent technical support, including Xenopus oocyte isolation and molecular biology; L. Bianchi for the gift of the UNC-8-encoding plasmid. This work was funded by fellowship to SF (DFG), the Amgen Scholars Program to ID (SSRP), and grants from NIH to MBG (R01NS07715, R35NS105092) and support from HHMI to LT.

---

**Supplementary-Figure 1:**
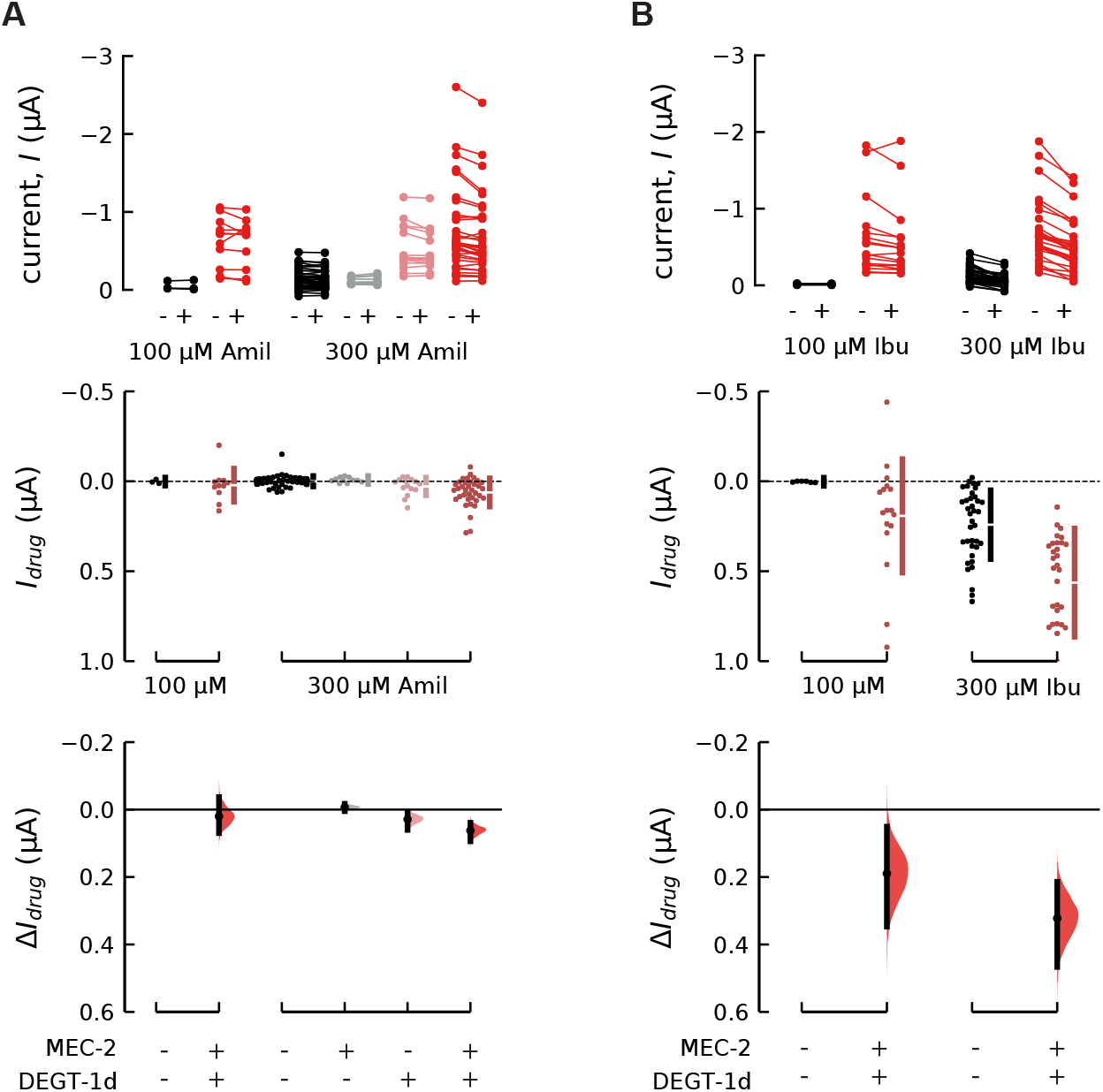
Sensitivity of DEGT-1d, DEGT-1 alone and MEC-2 alone to higher concentrations of amiloride and ibuprofen. **(A)** Top: Paired dots show the current, I at −85 mV in uninjected oocytes (black) and in oocytes expressing MEC-2 alone (grey), DEGT-1d alone (light red) and DEGT-1d (together with MEC-2) (red) before and after treatment with 100 μM or 300 μM Amil. Drug-sensitive current (I_*drug*_) (middle) at −85 mV in uninjected oocytes and oocytes expressing MEC-2 alone (grey), DEGT-1d alone (light red) and DEGT-1d (together with MEC-2) (red) compared to uninjected oocytes (black). Estimation plots (bottom) showing the effect of each drug (*ΔI*_*drug*_) on MEC-2 alone (grey), DEGT-1d alone (light red) and DEGT-1d (together with MEC-2) ( red) relative to the drug effect on uninjected oocytes. Subsequent estimation statistics are described as a change in current compared to the change in current to uninjected oocytes, the 95% confidence interval [in μA], and the two-sided p-value of the Mann-Whitney test: 100 μM Amil DEGT-1d: 0.111 μA [−0.00403, 0.388], p = 0.279 (n =12), 300 μM Amil DEGT-1d: 0.0623 μA [0.0383, 0.0922], 300 μM Amil DEGT-1d: 0.0623 μA [0.0383, 0.0922], p = 1.32e-06 (n = 36), DEGT-1 alone: 0.0288 μA [0.00847, 0.0617], p = 0.0528 (n = 15), MEC-2 alone: −0.0067 μA [−0.0176, 0.00437], p = 0.19 (n = 11). **(B)** Paired dots (top) show the current, *I* at −85 mV in uninjected oocytes (black) and in oocytes expressing DEGT-1d (red) before and after treatment with 100 μM or 300 μM Ibu. Drug-sensitive current (I*drug*) (middle) at −85 mV in oocytes expressing DEGT-1d (red) compared to uninjected oocytes (black). Estimation plots (bottom) showing the effect of each drug (*ΔI*_*drug*_) on DEGT-1d ( red) relative to the drug effect on uninjected oocytes. Subsequent estimation statistics are: 100 μM Ibu DEGT-1d: 0.19 μA [0.0446, 0.329], p = 0.0229 (n =17), 300 μM Ibu DEGT-1d: 0.322 [0.213, 0.465], p = 2.48e-06 (n = 30).

**Supplementary-Figure 2:**
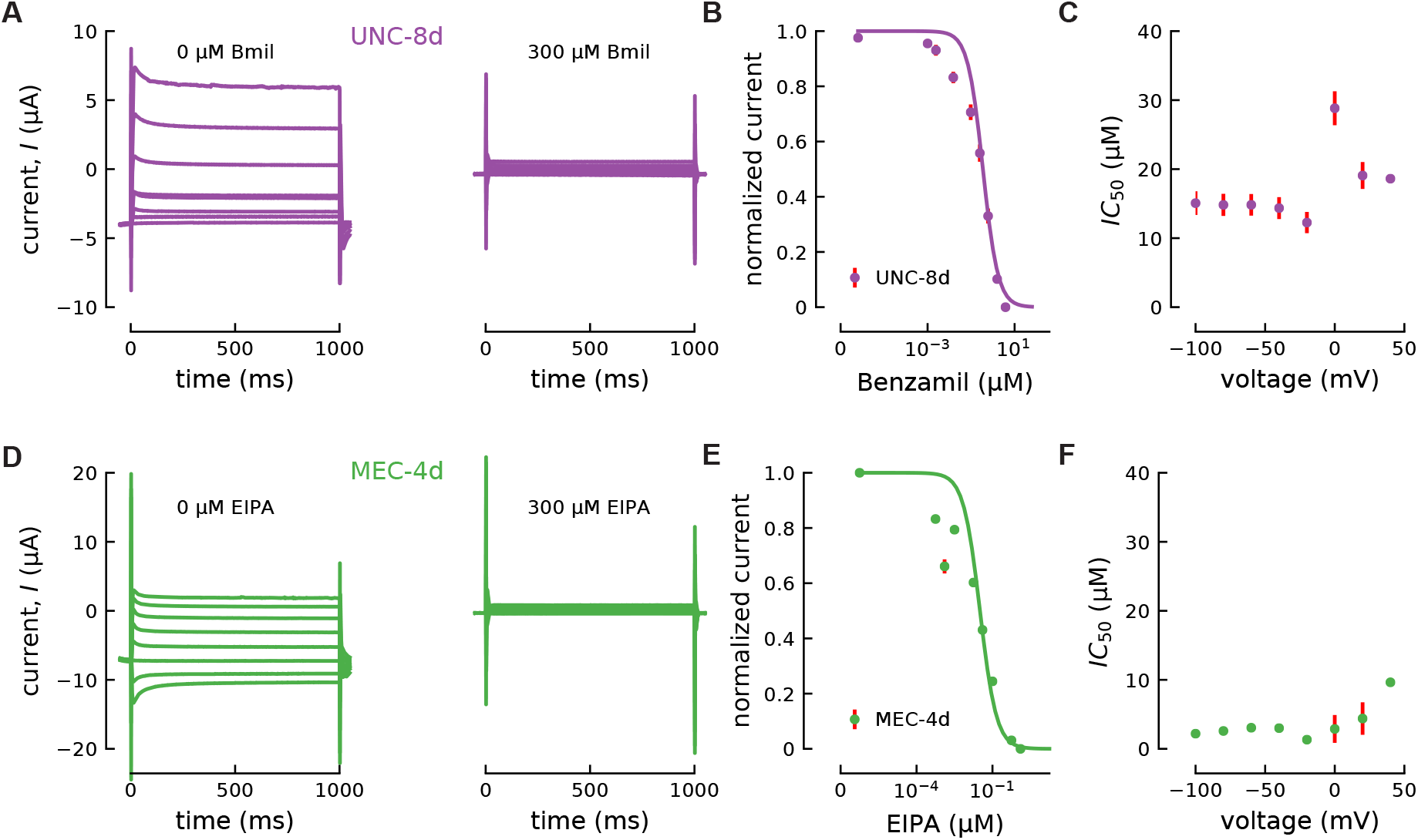
Sensitivity of UNC-8d channels to Benzamil and MEC-4d channels to EIPA. **(A)** Representative traces of currents of oocytes expressing UNC-8d channels in the absence (left) and presence (right) of 300 μM Benzamil (Bmil). Current responses to voltage steps from −100 mV to +40 mV in 20 mV increments. **(B)** Dose-response curves of UNC-8d channels to Bmil at −60 mV. The mean *IC*_50_ ± SEM for Bmil at −60 mV was 14.8 ± 1.6 μM (n = 4). **(C)***IC*_50_ values for different voltages (n = 4). Mean values for dose-response curves were derived from a step protocol similar to Fig. 1A. Instead of the ramp (red background), voltage steps from −100 mV to +40 mV in 20 mV increments were applied. Individual recordings were fitted with the Hill equation (*I*_*max*_ * *IC*_50_^*n*^/(*IC*_50_^*n*^ + *x*^*n*^)), where the Hill parameter was set to 1 and x is the concentration of the drug. **(D)** MEC-4d current in the absence (left) and presence (right) of 300 μM EIPA. Voltage pulses were applied between −100 to 40 mV (20 mV increment) from a holding potential of −60 mV, as shown in in Fig. 2A. Similar results were obtained in a total of 11 oocytes isolated from 3 frogs. **(E)** EIPA dose-response relationship of MEC-4d current at −60 mV (Normalized to *I*_max_ and baseline current). Points are the mean ± SEM of at least analyzed from at least 3 donor frogs and the smooth line is fit to these points using a single-binding site curve. **(F)** *IC*_50_ as a function of voltage (points are the mean±sem, n = 11, except 0 + 20 mV n=2 and +40 mV n =1).

